# IPMK modulates FFA-induced insulin resistance in primary mouse hepatocytes

**DOI:** 10.1101/2023.04.26.538310

**Authors:** Ik-Rak Jung, Rexford S. Ahima, Sangwon F. Kim

**Author notes:** **Corresponding Authors:** Sangwon F. Kim, PhD, Department of Medicine, Division of Endocrinology, Diabetes, and Metabolism, Johns Hopkins University, 5501 Hopkins Bayview Cir, AAC 2A44/62, Baltimore, Maryland, USA. 21224.

## Abstract

Insulin resistance is a critical mediator of the development of non-alcoholic fatty liver disease (NAFLD). An excess influx of fatty acids to the liver is thought to be a pathogenic cause of insulin resistance and the development of non-alcoholic fatty liver disease (NAFLD). Although elevated levels of free fatty acids (FFA) in plasma contribute to inducing insulin resistance and NAFLD, the molecular mechanism is not completely understood. This study aimed to determine whether inositol polyphosphate multikinase (IPMK), a regulator of insulin signaling, plays any role in FFA-induced insulin resistance in primary hepatocytes. Here, we show that excess FFA decreased IPMK expression, and blockade of IPMK decrease attenuated the FFA-induced suppression of Akt phosphorylation in primary mouse hepatocytes (PMH). Moreover, overexpression of IPMK prevented the FFA-induced suppression of Akt phosphorylation by insulin, while knockout of IPMK exacerbated insulin resistance in PMH. In addition, treatment with MG132, a proteasomal inhibitor, inhibits FFA-induced decrease in IPMK expression and Akt phosphorylation in PMH. Furthermore, treatment with the antioxidant N-Acetyl Cysteine (NAC) significantly attenuated the FFA-induced reduction of IPMK and restored FFA-induced insulin resistance in PMH. In conclusion, our findings suggest that excess FFA reduces IPMK expression and contributes to the FFA-induced decrease in Akt phosphorylation in PMH, leading to insulin resistance. Our study highlights IPMK as a potential therapeutic target for preventing insulin resistance and NAFLD.

## Introduction

IPMK is a key enzyme for inositol polyphosphate biosynthesis and plays a role in the metabolic signaling pathway by acting kinase independently via protein-protein interactions. The kinase-independent role of IPMK has been highlighted with findings that IPMK acts as an adapter protein to stabilize protein complexes regulated by energy status or nutrients. IPMK is a novel AMP–activated protein kinase (AMPK)-binding protein, and its interaction is regulated by glucose levels (Bang et al., 2012). Glucose deprivation reduces the IPMK-AMPK interaction, which is recovered by glucose supplements. In addition, IPMK mRNA and protein levels are suppressed in the hypothalamus during fasted states and are restored by refeeding (Bang et al., 2012). IPMK is also known to interact with mTOR (mammalian target of rapamycin), modulating mTOR complex 1 stability and amino acid-induced mTOR signaling (Kim et al., 2011). Loss of IPMK in murine embryonic fibroblasts (MEFs) reduces both amino acid-stimulated mTORC1 activation and cell growth (Kim et al., 2011). Mammalian IPMK contains a unique N-terminal region, which mediates its direct binding with mTOR. However, the regulation of IPMK by fatty acids or the role of IPMK in fatty acid-induced signaling is not well understood.

Free fatty acids (FFA) are the critical mediators of lipotoxicity and insulin resistance. An elevation of plasma FFA, as found in obesity with metabolic complications, such as Nonalcoholic fatty liver disease (NAFLD) and type 2 diabetes mellitus (T2DM), induces hepatic insulin resistance (Biden, Boslem, Chu, & Sue, 2014; Boden et al., 2005; Pan et al., 2019; Pereira et al., 2013). Lipid overload in the liver promotes mitochondrial β-oxidation resulting in disruption of fatty acid metabolism and excessive electron flux in the mitochondria respiratory chain, which can cause ROS generation and impaired insulin signaling pathway (Liu et al., 2017). Furthermore, oxidative stress has been implicated as one of the key mechanisms in the pathogenesis of insulin resistance. It has been reported that in vitro models of insulin resistance in cultured adipocytes and myotubes are associated with mitochondrial oxidative stress (Fazakerley et al., 2018; Hoehn et al., 2009). Enhanced ROS production can affect various points in insulin receptor and intracellular signaling, such as JNK, p38, and IKKβ, ultimately causing insulin resistance (Tiganis, 2011). Also, oxidative stress has been reported as one of the main contributors to FFA-induced hepatic insulin resistance (Nakamura et al., 2009).

Recently, we have shown that IPMK plays an essential role in hepatic insulin signaling and whole-body glucose metabolism (Jung, Anokye-Danso, Jin, Ahima, & Kim, 2022). Our group reported that the expression of IPMK is decreased in the liver of HFD-fed mice, and loss of hepatic IPMK exacerbates glucose and insulin tolerance in HFD-fed mice compared to WT mice (Jung et al., 2022), indicating that IPMK is involved in the pathogenesis of Insulin resistance. The current work aimed to determine whether a change in IPMK contributes to FFA-induced insulin resistance in the primary hepatocytes. We found that FFA treatment reduced IPMK expression in the primary hepatocytes, which was associated with decreased insulin-mediated activation of the Akt pathway. Importantly, inhibition of IPMK decrease in FFA-treated hepatocytes attenuates FFA-mediated insulin resistance, measured by Akt phosphorylation levels.

## Results

### FFA treatment reduces IPMK protein levels in primary mouse hepatocytes

It has been shown that IPMK regulates insulin signaling in multiple cell types. We recently reported that the deletion of hepatic IPMK decreased insulin-mediated Akt signaling and aggravated HFD-induced insulin resistance and glucose intolerance (Jung et al., 2022). Moreover, we found a significant decrease in IPMK protein in the liver of insulin-resistant mice subjected to a High-fat diet (HFD) for a long-term period (Jung et al., 2022). To examine whether a reduction in hepatic IPMK by HFD contributed to the induction of insulin resistance in mice, we investigated the effect of FFA on IPMK protein in primary mouse hepatocytes (PMH). PMH were treated with different concentrations of the FFA mixture (palmitic acid (PA): oleic acid (OA) in the molar ratio 1:2). FFA treatment decreased IPMK protein in a concentration-dependent manner (Fig. 1A). Treatment with 0.9 mM FFA was sufficient to reduce the IPMK protein, and 1.2 and 1.5 mM FFA enhanced to decrease IPMK protein. We also observed a time-dependent reduction in the protein level of IPMK by FFA treatment (Fig 1B). IPMK protein was significantly reduced at 8 hr after FFA treatment (Fig. 1B).

**Figure 1.**
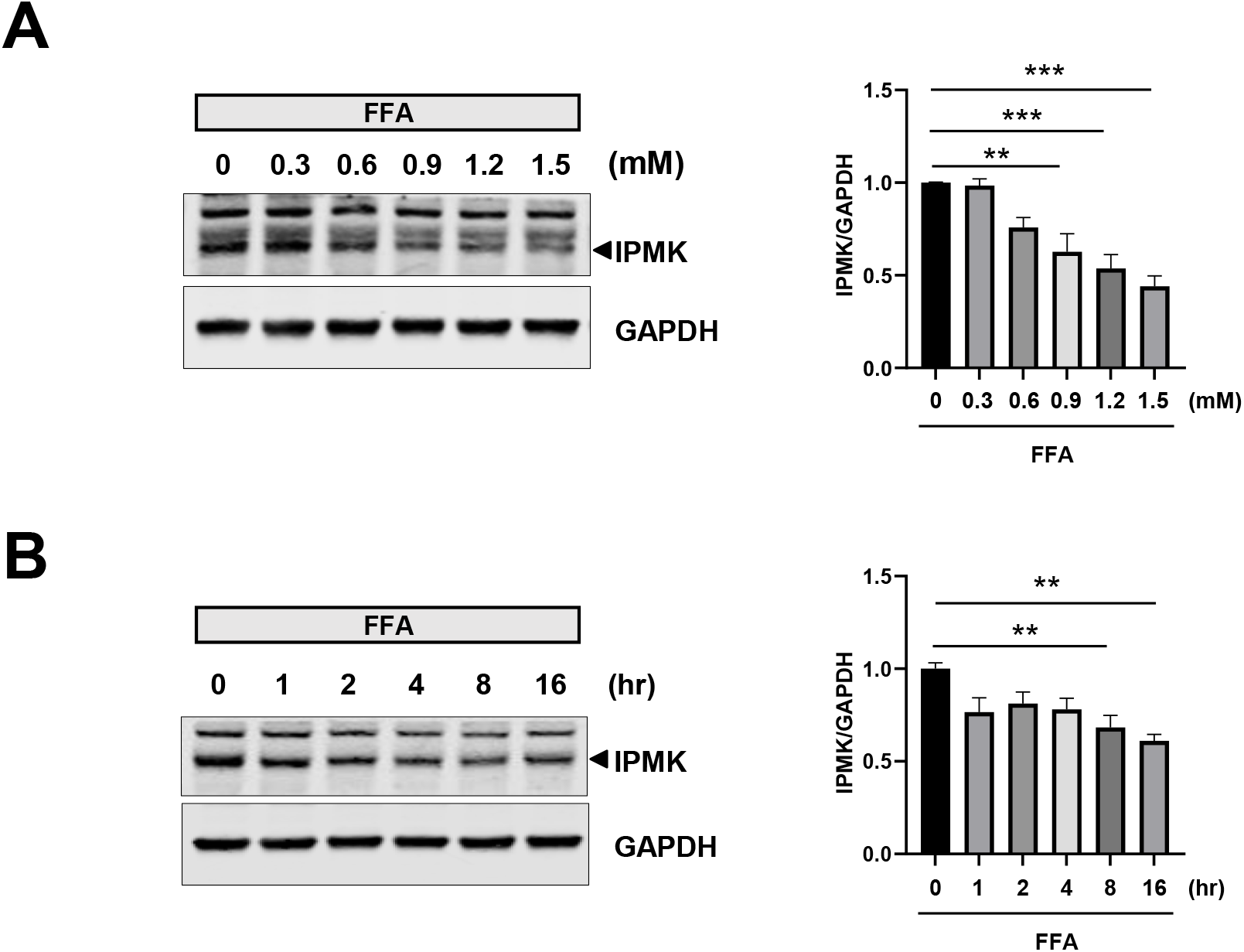
Reduction of IPMK protein in PMH by FFA treatment. (A) Primary mouse hepatocytes (PMH) were treated with different concentrations of FFA for 16 hr, and cell lysates were subjected to immunoblotting (B) PMH were treated with 0.9 mM FFA for various time points. The levels of IPMK expression were measured by immunoblotting. Levels of indicated proteins (IPMK vs. GAPDH) were quantified by LI-COR Image Studio software. The representative Western bolt images from at least 3 independent assays are presented. Data are presented as mean ± SEM; **p < 0.01, ***p < 0.001, n = 3.

### IPMK plays a role in FFA-induced insulin resistance in primary mouse hepatocytes

Since FFA reduced IPMK protein levels, we investigated whether a decrease in IPMK protein by FFA treatment is associated with insulin resistance observed in FFA-treated PMH. First, we confirmed that treatment with FFA for 16 hr significantly reduced IPMK protein levels and decreased insulin-stimulated Akt phosphorylation at both T308 and S473 at the same time (Fig. 2). Next, we examined the effects of IPMK overexpression on insulin-mediated phosphorylation of Akt in FFA-treated PMH. FFA reduced endogenous IPMK protein level and insulin-stimulated Akt phosphorylation in PMH, while overexpression of IPMK significantly attenuated FFA-induced reduction of Akt phosphorylation in PMH (Fig 3A). To better understand the effects of IPMK levels on FFA-induced insulin resistance in PHM, we also investigated whether the knockdown of IPMK could aggravate FFA-induced reduction of Akt phosphorylation in PMH. We isolated PMH from IPMK loxp mice (Maag et al., 2011) and treated them with an adeno-Cre virus (Ad-Cre) to knockdown IPMK. We confirmed that IPMK protein was almost completely decreased by Ad-Cre treatment in PMH (Fig 3B). We found that a loss of IPMK in PMH appeared to exacerbate FFA-mediated insulin resistance without statistical significance (Fig 3B). Overall, our data suggest that the FFA-induced reduction of IPMK is associated with a decrease in Akt phosphorylation in PMH treated with FFA.

**Figure 2.**
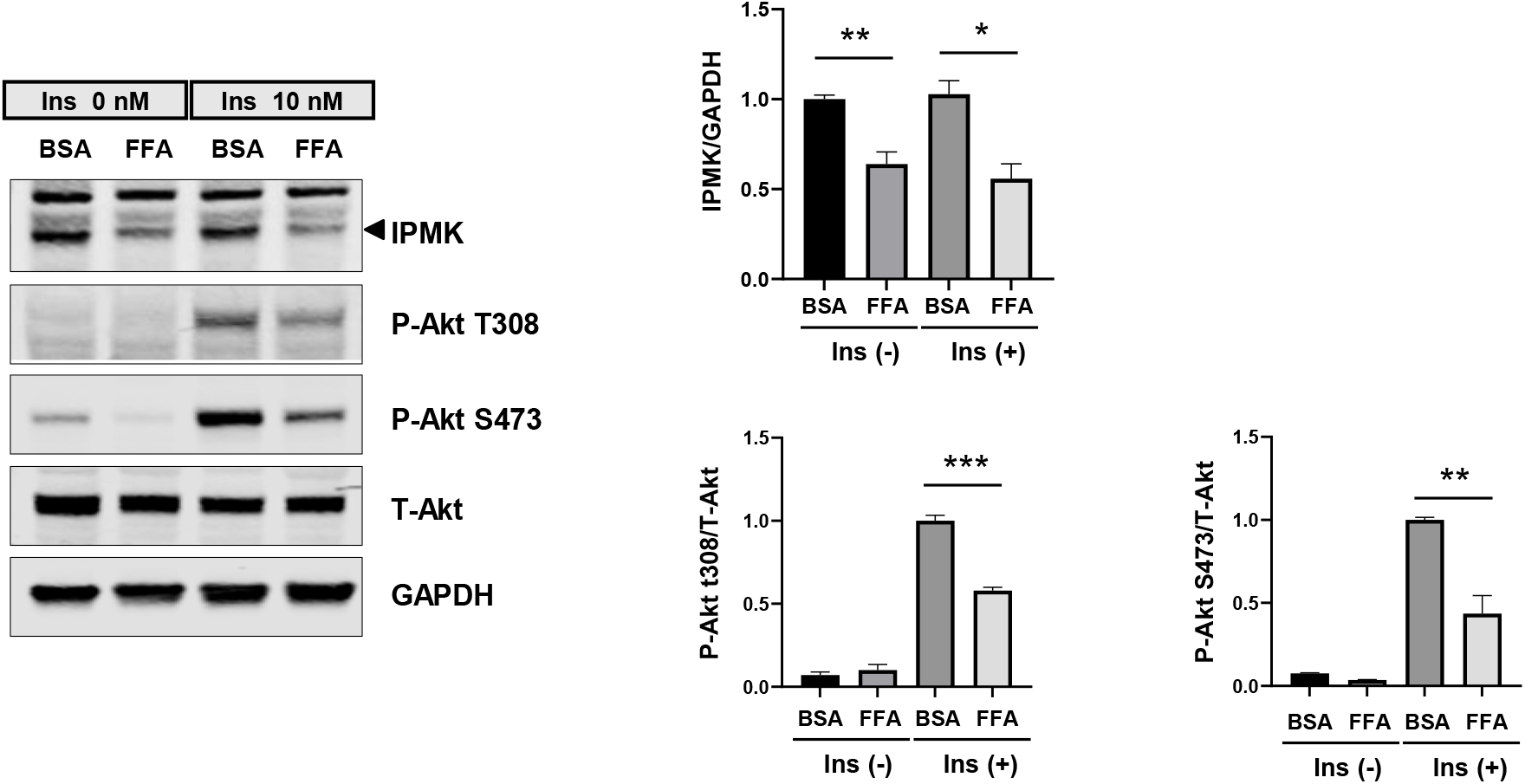
Effect of p-Akt in PMH by FFA treatment. Insulin resistance by FFA treatment was determined by measuring levels of phospho-protein kinase B (p-Akt) in immunoblotting after 10 min insulin stimulation (10 nM). Levels of indicated proteins (phosphorylated vs. total Akt or IPMK vs. GAPDH) were quantified by LI-COR Image Studio software. The representative Western bolt images from at least 3 independent assays are presented. Data are presented as mean ± SEM; *p < 0.05, **p < 0.01, ***p < 0.001, n = 3.

**Figure 3.**
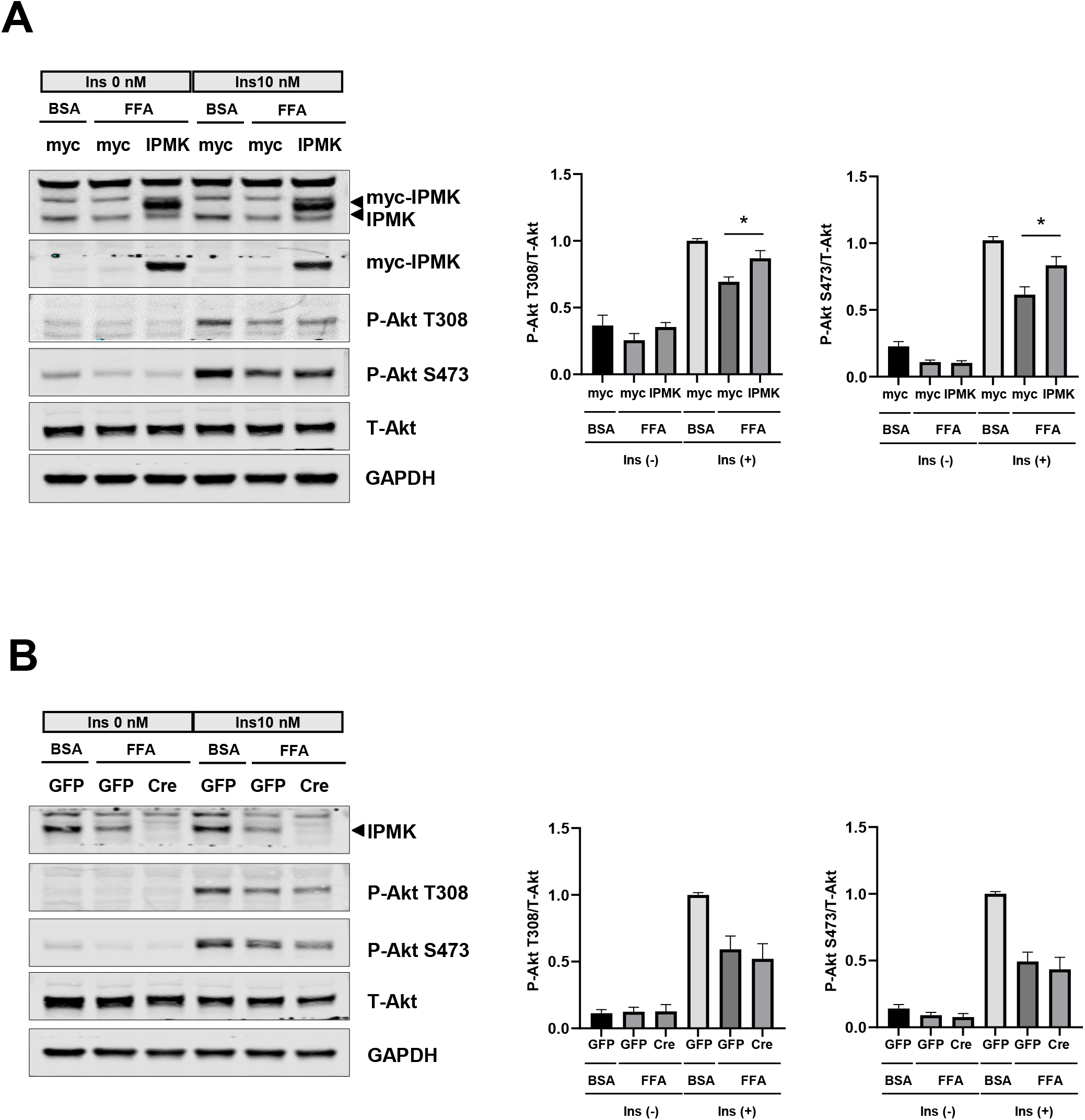
Protective effect of IPMK overexpression on FFA-induced insulin resistance in PMH. (A) PMH were overexpressed with IPMK, and insulin resistance by FFA treatment was determined by measuring levels of phospho-Akt in immunoblotting after 10 min insulin stimulation (10 nM). (B) PMH were treated with Adenovirus-GFP (Ad-GFP) or Adenovirus-Cre (Ad-Cre), and insulin resistance by FFA treatment was determined by measuring levels of phospho-Akt in immunoblotting after 10 min insulin stimulation (10 nM). Levels of indicated proteins (phosphorylated vs. total Akt) were quantified by LI-COR Image Studio software. The representative Western bolt images from at least 3 independent assays are presented. Data are presented as mean ± SEM; *p < 0.05, n = 3.

### Inhibition of IPMK degradation improves FFA-induced insulin resistance in PMH

We found that FFA treatment led to a decrease in IPMK expression and suppression of insulin-mediated phosphorylation of Akt, yet overexpression of IPMK attenuated FFA-induced reduction of Akt phosphorylation in PMH; hence we wondered whether prevention of IPMK degradation could protect against FFA-induced decrease of Akt phosphorylation in PMH. MG132, a proteasome inhibitor, is commonly used to prevent proteasome activity (Kisselev & Goldberg, 2001). To investigate whether MG132 prevented the FFA-induced IPMK protein degradation, PMH were treated with FFA in the presence of MG132 for 16 hr. FFA-induced IPMK protein degradation was blocked by MG132 treatment (Fig. 4). Importantly, FFA-induced reduction of Akt phosphorylation at T308 was significantly prevented by MG132 treatment (Fig. 4). These results suggest that FFA-induced IPMK reduction is partly due to proteasomal degradation and plays a role in FFA-induced decrease in Akt phosphorylation in PMH.

**Figure 4.**
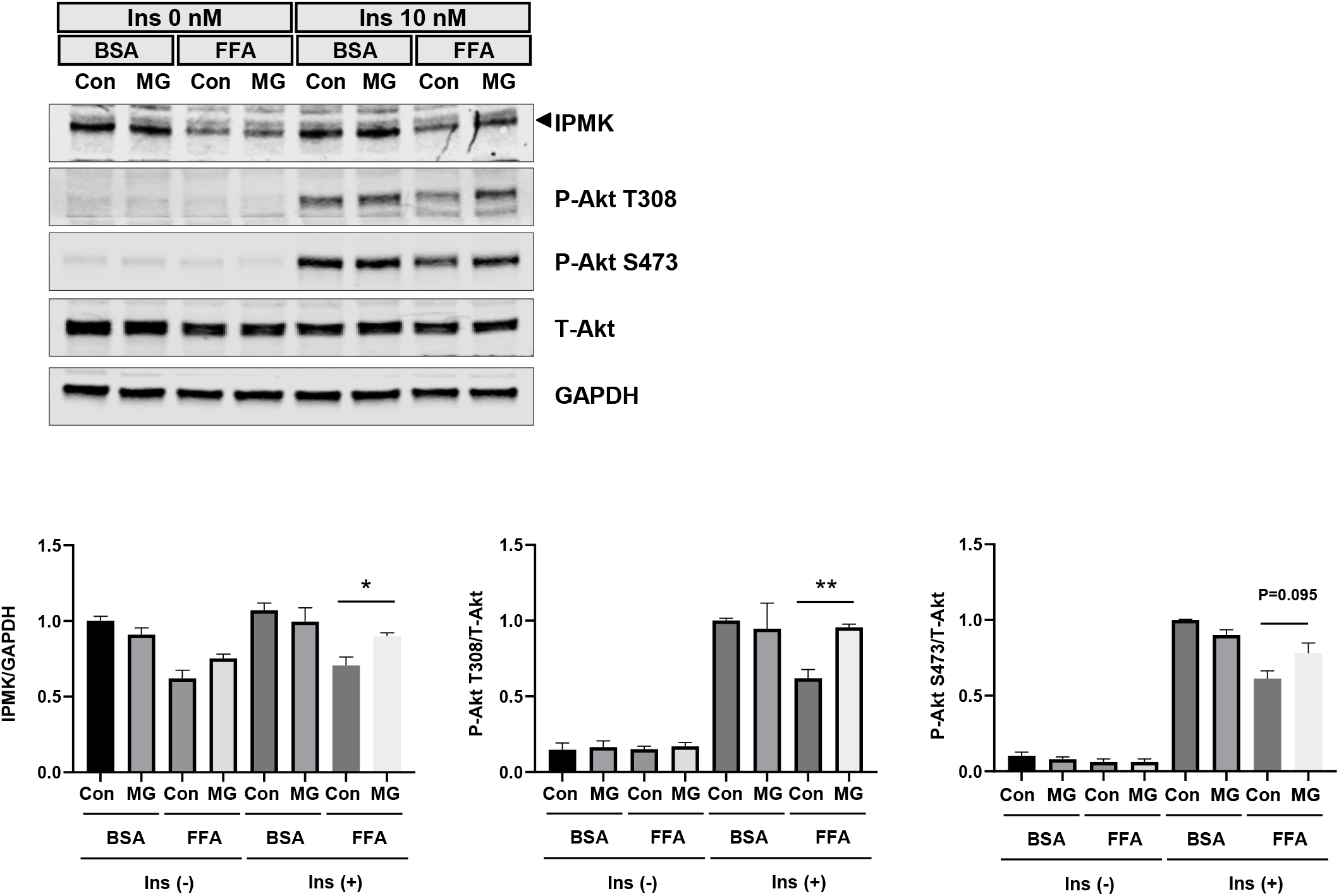
Involvement of proteasomal degradation in FFA-induced IPMK reduction. PMH were treated with 0.9 mM FFA in the presence of 1 uM MG132 for 16 hr. The protective effect of IPMK reduction and Insulin resistance by FFA treatment was determined by measuring levels of IPMK and phospho-protein kinase B (p-Akt) in immunoblotting after 10 min insulin stimulation (10 nM). Levels of indicated proteins (phosphorylated vs. total Akt or IPMK vs. GAPDH) were quantified by LI-COR Image Studio software. The representative Western bolt images from at least 3 independent assays are presented. Data are presented as mean ± SEM; *p < 0.05, **p < 0.01, n = 3.

### Antioxidant attenuates IPMK degradation and improves insulin resistance in FFA-treated PMH

The elevated influx of FFA exceeding the metabolic capacity of hepatocytes causes incomplete beta-oxidation and mitochondrial dysfunction (Clare, Dillon, & Brennan, 2022), which leads to ROS and oxidative stress, ultimately resulting in insulin resistance (Hurrle & Hsu, 2017). ROS are associated with metabolic processes, and oxidized proteins are more efficiently degraded (Forrester, Kikuchi, Hernandes, Xu, & Griendling, 2018; Raynes, Pomatto, & Davies, 2016). N-Acetyl Cysteine (NAC), a cysteine prodrug, is widely used as a pharmacological antioxidant (Ezeriņa, Takano, Hanaoka, Urano, & Dick, 2018). To determine whether NAC treatment could prevent FFA-induced IPMK reduction, we treated FFA and NAC to PMH for 16 hr. FFA-induced IPMK protein degradation was significantly prevented by NAC treatment (Fig. 5). Similar to MG132 treatment, FFA-induced reduction of Akt phosphorylation at T308 was significantly blocked by NAC treatment (Fig. 5). These results suggest that FFA-induced IPMK reduction is at least in part due to ROS production and scavenging ROS could attenuate FFA-induced Akt phosphorylation reduction.

**Figure 5.**
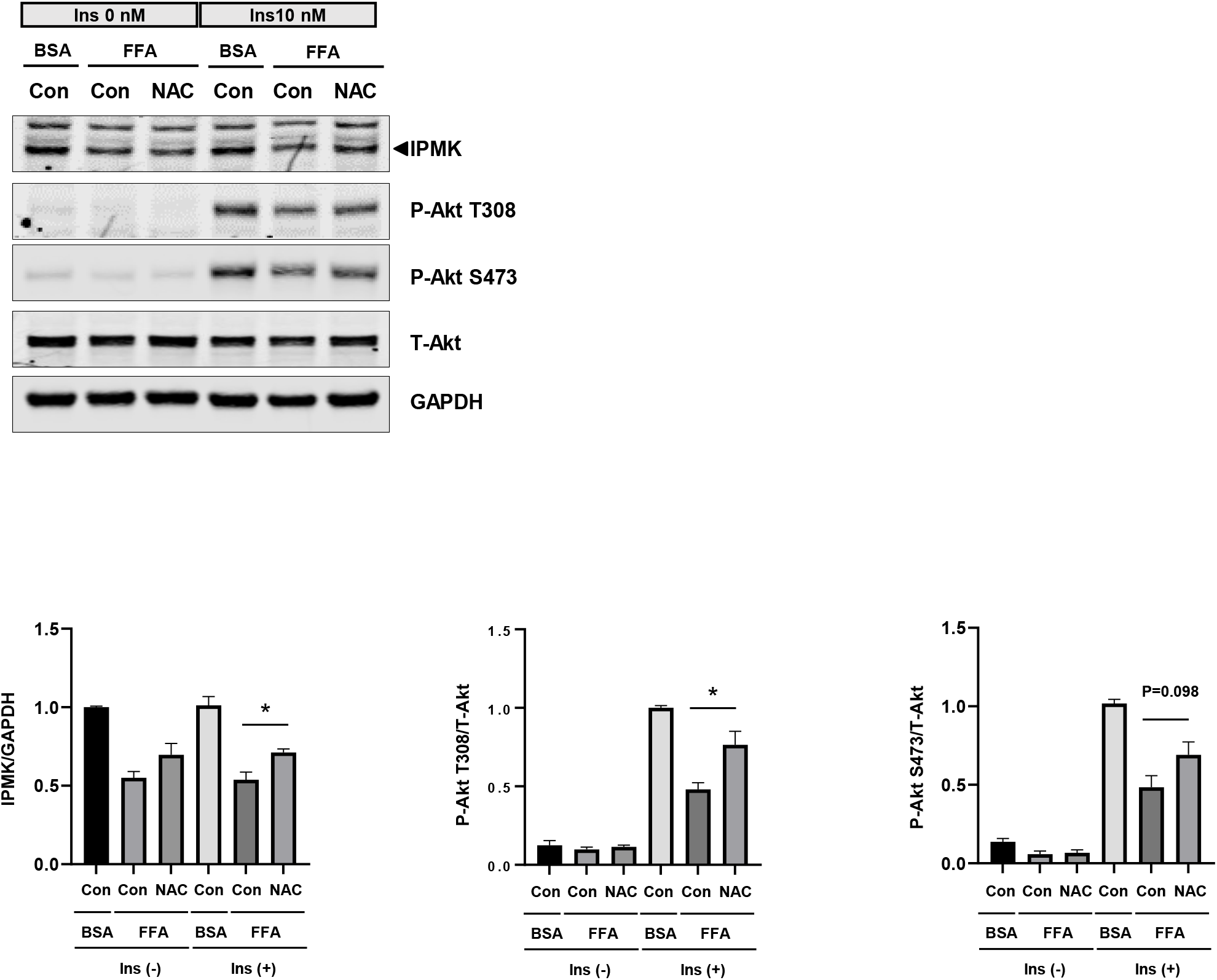
Effect of antioxidant on FFA-induced IPMK reduction. PMH were treated with 0.9 mM FFA in the presence of 1 uM NAC for 16 hr. The protective effect of IPMK reduction and Insulin resistance by FFA treatment was determined by measuring levels of IPMK and phospho-protein kinase B (p-Akt) in immunoblotting after 10 min insulin stimulation (10 nM). Levels of indicated proteins (phosphorylated vs. total Akt or IPMK vs. GAPDH) were quantified by LI-COR Image Studio software. The representative Western bolt images from at least 3 independent assays are presented. Data are presented as mean ± SEM; *p < 0.05, n = 3.

## Discussion

As an energy and nutrient sensing protein, IPMK is regulated by glucose and amino acids and is reported to be involved in these signals (Bang et al., 2012; Kim et al., 2011). However, neither the regulation of IPMK by lipids nor the role of IPMK in lipid metabolism has been reported. In this study, we investigated the hypothesis that IPMK plays a role in HFD-induced insulin resistance. IPMK protein levels were decreased in the liver tissues of mice fed HFD (Jung et al., 2022). Moreover, FFA treatment decreased IPMK protein in PMH in a concentration- and time-dependent manner. A decrease of IPMK protein by FFA was significantly prevented by a proteasomal inhibitor or antioxidant treatment, accompanied by the recovery of insulin signaling in PMH. Furthermore, IPMK overexpression restored the FFA-induced decrease in Akt phosphorylation, whereas knockout of IPMK tended to exacerbate it in PMH. Together, these data suggest that the reduction of IPMK by excess FFA plays a role in FFA-induced suppression of Akt phosphorylation in PMH and that hepatic IPMK may be of therapeutic target in preventing diet-induced insulin resistance.

The elevated FFA levels are closely correlated to insulin resistance in obese patients with type 2 diabetes (Boden, 1999; Spiller, Blüher, & Hoffmann, 2018). In rodents, i.v. Infusions of fatty acids increased plasma FFA and induced hepatic insulin resistance (Pereira et al., 2021). It has been suggested that FFA causes insulin resistance by the formation of lipid intermediates such as diacylglycerol (Erion & Shulman, 2010), lysophosphatidic acid (Nagle et al., 2007), ceramides (Sokolowska & Blachnio-Zabielska, 2019), cellular stress including oxidative stress and endoplasmic reticulum (ER) stress (Eizirik, Cardozo, & Cnop, 2008; Tangvarasittichai, 2015) and proinflammatory cytokines (e.g., TNF-a, IL-1b, IL-6, and MCP1) (Shoelson, Herrero, & Naaz, 2007). Despite numerous studies investigating the potential mechanism of FFA-induced insulin resistance, it is still not completely understood. Our study found that IPMK protein was significantly decreased in PMH treated with FFA. It is known that IPMK can regulate insulin signaling in multiple cell types (Arnold et al., 2014; Guha et al., 2020; Maag et al., 2011). Furthermore, we recently have demonstrated that IPMK protein is decreased in the liver of mice fed an HFD and that deletion of hepatic IPMK suppressed insulin-mediated Akt signaling and exacerbated HFD-induced insulin resistance (Jung et al., 2022), suggesting that a decrease in IPMK protein may be partly involved in HFD- and FFA-induced insulin resistance in vivo and in vitro, respectively.

Although several studies have reported that IPMK plays a role in regulating insulin signaling, how IPMK is regulated is yet to be examined. Bang et al. reported that glucose deprivation downregulates IPMK mRNA and protein levels in MEFs. IPMK mRNA and protein levels in the hypothalamus are significantly decreased in fasted mice (Bang et al., 2012). Recently, Lee et al. reported that both mRNA and protein levels of IPMK were decreased in mouse and human macrophages by lipopolysaccharide (LPS) (Lee, Kim, & Kim, 2023). However, no studies have investigated the IPMK levels in metabolic dysfunction. We reported a significant decrease in IPMK protein in the fatty liver subjected to HFD for a long-term period (Jung et al., 2022). To the best of our knowledge, our finding is the first to demonstrate a regulation of IPMK protein in metabolic dysfunction potentially contributing to insulin resistance. PA and OA are the most abundant FFAs in liver TG in a patient with NAFLD (Aoudjehane et al., 2020). PA/OA-treatment increased lipid accumulation with decreased insulin signal in PMH (Choi et al., 2022). To investigate the mechanism of how HFD feeding downregulates IPMK protein in the liver, we used the model of PA/OA (FFA)-induced steatosis in PMH to mimic the HFD-induced steatosis in mice (Soret, Magusto, Housset, & Gautheron, 2020). Indeed, we showed that FFA treatment significantly decreases IPMK protein in PMH in a dose- and time-dependent manner (Fig.1A and B). Saturated fatty acid such as palmitic acid has been shown to upregulate the proteolytic system in skeletal muscle (Perry et al., 2018; Woodworth-Hobbs et al., 2014; Yang et al., 2013), and facilitates ubiquitination of cellular proteins leading to their degradation via the ubiquitin– proteasome pathway in HepG2 cells (Ishii, Maeda, Tani, & Akagawa, 2015). It is also reported that PA treatment led to a reduction in heterogeneous nuclear ribonucleoproteins (hnRNPs) and serine-rich splicing factor 3 (SRSF3) protein levels by neddylation-mediated proteasomal degradation in AR42J and HepG2 cells, respectively (Chen, Wang, Xia, Zhang, & Zhao, 2022; Kumar et al., 2019). Since MG132 significantly prevented FFA-induced IPMK reduction, we speculate that FFA-mediated modification of IPMK may trigger its degradation via proteasomal systems.

Mitochondrial dysfunction has been considered pathogenically linked to insulin resistance (Choi et al., 2022) and suggested that excessive FFA into mitochondria leads to incomplete FA oxidation, producing an increased generation of reactive oxygen species (ROS), ultimately resulting in cellular damage and insulin resistance through oxidative stress (Palomer, Pizarro-Delgado, Barroso, & Vazquez-Carrera, 2018). In particular, ROS can damage macromolecules, including proteins. When proteins are irreversibly damaged by ROS, cells require systems for recognition and elimination, such as the lysosomal system (Settembre, Fraldi, Medina, & Ballabio, 2013), mitochondrial proteases (Ngo, Pomatto, & Davies, 2013), and proteasome system (Grimm, Höhn, & Grune, 2012). Our study demonstrated that not only a proteasomal inhibitor but also antioxidant treatment prevented IPMK protein from excessive FFA, suggesting that FFA-induced oxidative stress and proteasomal degradation may modulate the levels of IPMK protein. Thus, further studies are required to elucidate how IPMK levels are regulated by FFA in hepatocytes. In conclusion, we have demonstrated that excessive FFA decreases hepatic IPMK, and preventing reduction of IPMK restores FFA-induced decrease of Akt phosphorylation in PMH. Attempts to prevent IPMK degradation or maintain IPMK protein may provide a protective strategy for improving insulin resistance in obesity with type 2 diabetes.

## Materials and Methods

### Cell Culture

Primary hepatocytes were isolated from WT male mice, using collagenase perfusion followed by filtration through a 70 μm mesh as previously described (Han et al., 2016). Cells were all maintained in Media 199 (Corning Inc, Corning, NY, USA) supplemented with 10% fetal bovine serum (MilliporeSigma, Burlington, MA, USA) and 100 units/mL penicillin/streptomycin (MilliporeSigma, Burlington, MA, USA). Primary hepatocyte transfection was performed using a lipofectamine 3000 transfection reagent (Thermo Fisher Scientific, Waltham, MA, USA), according to the manufacturer’s protocol.

### Preparation of FFAs

Palmitate/oleate/bovine serum albumin (BSA) conjugates were prepared by soaping palmitate or oleate with sodium hydroxide and mixing with BSA. Palmitate or oleate (20 mM in 0.01 M NaOH) was incubated at 70 C for 30 minutes. Fatty acid soaps were then complexed with 5% fatty acid-free BSA in phosphate-buffered saline (PBS) at a 1: 3 volume ratios. Complexed fatty acids consisted of 5 mM palmitate or oleate and 3.75% BSA. The palmitate/oleate/BSA conjugates were diluted in Media199 medium containing 10 % FBS and administered to cultured cells.

### Immunoblotting

Samples were lysed in RIPA buffer containing inhibitors and heated at 95°C for 5 min before electrophoresis. Proteins were transferred to a 0.2-mm nitrocellulose membrane, blocked with 5% nonfat dry milk, and incubated with primary antibodies at 4°C overnight. Immunoblotting was conducted with the following antibodies: IPMK from Covance, p-Akt-S473, p-Akt-T308, t-Akt, and GAPDH from Cell Signaling Technology. Blots were imaged and quantitated using an Odyssey Near-Infrared Scanner (Li-Cor Biosciences, Lincoln, NE, USA).

### Statistical Analysis

All statistical analyses were performed using GraphPadPrism 5.0 (GraphPad, San Diego, CA, USA), and all data are presented as mean ± SEM. Comparisons between the two groups were made using Student’s t-test. Comparisons among multiple groups at a one-time point were made using one-way ANOVA. The threshold for statistical significance was set at P < 0.05, and Bonferroni’s multiple-comparison test was used for all post hoc analyses.

## Author contributions

SFK and IJ designed and directed the project. IJ performed the experiment. IJ analyzed data and wrote the first draft of the manuscript. RSA and SFK reviewed and edited the manuscript. All authors approved the paper for submission.

## Acknowledgments

This work was supported by grants from American Heart Association Strategically Funded Research Network (SFRN) Obesity Center (RSA and SFK, 17SFRN33610014 and 20SFRN35210662), National Institutes of Health/National Institute of Diabetes and Digestive and Kidney Diseases (NIH/NIDDK) (SFK and RSA, DK135751), and American Heart Association SFRN fellowship (IJ, 17SFRN33560006).

